# Polyester: Simulating RNA-Seq Datasets With Differential Transcript Expression

**DOI:** 10.1101/006015

**Authors:** Alyssa C. Frazee, Andrew E. Jaffe, Ben Langmead, Jeffrey T. Leek

## Abstract

**Motivation:** Statistical methods development for differential expression analysis of RNA sequencing (RNA-seq) requires software tools to assess accuracy and error rate control. Since true differential expression status is often unknown in experimental datasets, artificially-constructed datasets must be utilized, either by generating costly spike-in experiments or by simulating RNA-seq data.

**Results:** *Polyester* is an R package designed to simulate RNA-seq data, beginning with an experimental design and ending with collections of RNA-seq reads. Its main advantage is the ability to simulate reads indicating isoform-level differential expression across biological replicates for a variety of experimental designs. Data generated by Polyester is a reasonable approximation to real RNA-seq data and standard differential expression workflows can recover differential expression set in the simulation by the user.

**Availability and Implementation:** *Polyester* is freely available from Bioconductor (http://bioconductor.org/).

**Supplementary Information:** Supplementary figures are available online.

## 1 Introduction

RNA sequencing (RNA-seq) experiments have become increasingly popular as a means to study gene expression. There are a range of statistical methods for differential expression analysis of RNA-seq data (Oshlack *et al.*, 2010). The developers of statistical methodology for RNA-seq need to test whether their tools are performing correctly. Often, accuracy tests cannot be performed on real datasets because true gene expression levels and expression differences between populations are usually unknown, and spike-in experiments are costly in terms of both time and money.

Instead, researchers often use computational simulations to create datasets with a known signal and noise structure. Typically, simulated expression measurements used to evaluate differential expression tools are generated as gene counts from a statistical model like those used in common differential expression tools(Robinson *et al.*,2010; Anders and Huber, 2010). But these simulated scenarios do not account for variability in expression measurements that arises during upstream steps in RNA-seq data analysis, such as read alignment or read counting. *Polyester* is a new R package for simulating RNA-seq reads. *Polyester*’s main advantage is that users can simulate sequencing reads with specified differentialexpression signal for either genes or isoforms. This allows users to investigate sources of variability at multiple points in RNA-seq pipelines.

Existing RNA-seq simulators that generate sequencing reads are not designed for simulating experiments with biological replicates and specified differential expression signal. For example, the `rsem-simulate-reads` utility shipped with RSEM (Li and Dewey, 2011) requires a time-consuming first step of aligning real sequencing reads to developa sequencing model before reads can be simulated, and differential expression simulation is not built-in. Neither FluxSimulator (Griebel *et al.*, 2012) norBEERS (Grant *et al.*, 2011) have a built-in mechanism for introducing differential expression. These simulators also do not provide methods for defining a model for biological variability across replicates or specifying the exact expression level of specific transcripts. TuxSim has been used to simulate RNA-seq datasets with differential expression (Trapnell *et al.*, 2013), but it is not publicly available.

*Polyester* was created to fulfill the need for a tool to simulate RNA-seq reads for an experiment with replicates and well-defined differential expression. Users can easily simulate small experiments from a few genes or a single chromosomeThis can reduce computational time in simulation studies when computationally intensive steps such as read alignment must be performed as part of the simulation. *Polyester* is open-source, cross-platform, and freely available for download at https://github.com/alyssafrazee/polyester.

## 2 Methods

### 2.1 Input

*Polyester* takes annotated transcript nucleotide sequences as input.These can be provided as cDNA sequences in FASTA format, labeled by transcript. Alternatively, users can simulate from a GTF file denoting exon, transcript, and gene structure paired with full-chromosome DNA sequences. The flexibility of this input makes it possible to design small, manageable simulations by simply passing *Polyester* a FASTA or GTF file consisting of feature sets of different sizes. Efficient functions for reading, subsetting, and writing FASTA files are available in the *Biostrings* package (Pages *et al.*, ????), which is a dependency of *Polyester*.

### 2.2 RNA-Seq Data as Basis for Model Parameters

Several components of *Polyester*, described later in this section, require parameters estimated from RNA-seq data. To get these parameter estimates, we analyzed RNA-seq reads from 7 biological replicates in the public GEUVADIS RNA-seq data set (AC’t Hoen *et al.*, 2013; Lappalainen *et al.*,2013). The 7 replicates were chosen by randomly selecting one replicate from each of the 7 laboratories that sequenced samples in the study. These replicates represented 7 people from three different HapMap populations: CEU (Utah residents with Northern and Western European ancestry), TSI (Tuscani living in Italy), and YRI (Yoruba living in Ibadan, Nigeria). Data from the GEUVADIS study is available from the ArrayExpress database (http://www.ebi.ac.uk/arrayexpress) under accession numbers E-GEUV-1 through E-GEUV-6. We specifically used TopHat read alignments for these 7 replicates, under accession number E-GEUV-6. The reads were 75bp, paired-end reads.

Also available for the GEUVADIS data set is a fully processed transcriptome assembly, created based on the RNA-seq reads from all 667 replicates in the GEUVADIS study without using a reference transcriptome. This assembly was built using Cufflinks and processed with the *Ballgown* R package (Frazee *et al.*, ????), and is available for direct download as an R object (Leek, 2014).

### 2.3 Expression Models

A key feature of *Polyester* is that the analyst has full control over the number of reads that are generated from each transcript in the input file, for each replicate in the experiment. *Polyester* ships with a built-in model for these read numbers, or the model can be explicitly specified by the end user.

#### 2.3.1 Built-in Negative Binomial Read Count Model

The built-in transcript read count model assumes that the number of reads to simulate from each transcript is drawn from the negative binomial distribution, across biological replicates. The negative binomial model for read counts has been shown to satisfactorily capture biological and technical variability (Anders and Huber, 2010; Robinson *et al.*, 2010). In *Polyester*, differential expression between experimental groups is defined by a multiplicative change in the mean of the negative binomial distribution generating the read counts.

Specifically, define *Y_ijk_* as the number of reads simulated from replicate *i*, experimental condition *j*, and transcript *k*(*i* = 1, …,*n_j_*; *j* = 1,…,*J*; and *k* = 1,…, *N*; where *n_j_*is the number of replicates in condition *j*, *J* is the total number of conditions, and *N* is the total number of transcripts provided). The built-in model in *Polyester* assumes: *Y_ijk_* ~ Negative Binomial(mean = *μjk*, size = *r_jk_*)

In this negative binomial parameterization, *E*(*Y_ijk_*) = *μ _jk_* and Var 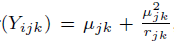 so each transcript’s expression variance across biological replicates is quadratically related to its baseline mean expression.The quantity 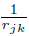 is commonly referred to as the dispersion parameter in this parameterization (Robinson *et al.*, 2010; Lawless, 1987; Ismail and Jemain, 2007). The user can provide *μ_jk_* for each transcript *k* and experimental group *j*. In particular, the user can relate transcript *k*’s length to *μ_jk_*. Also, this flexible parameterization reduces to the Poisson distribution as *r_jk_* → ∞. Since the Poisson distribution is suitable for capturing read count variability across technical replicates (Bullard *et al.*, 2010), users can create experiments with simulated technical replicates only by making all *r_jk_* very large. By default, 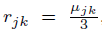, which means Var(*Y_jk_*) = 4*μ_jk_*.The user can adjust *r_jk_* on a per-transcript basis as needed, to explore different mean/variance expression models.

When *J* = 2, differential expression is set by providing a fold change λ between the two conditions for each transcript. Initially, a baseline mean *μ_k_* is provided for each transcript, and *μ_1k_*and *μ_2k_* are set to *μ_k_*. Then, if fold change λ is provided, *μ_1k_* and *μ_2k_* are adjusted:if λ > 1,*μ_1k_*, λ*μ_k_*and if λ < 1, 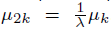 The number of reads to generate from each transcript is then drawn from the corresponding negative binomial distribution. When *J* > 2, the count for each transcript, *y_ijk_*, is generated from a negative binomial distribution with overall mean μ_k_ and size *r_jk_*. Differential expression can be set using a fold change matrix with *N* rows and *J* columns. Each count *y_ijk_* is multiplied by entry *k*, *j* of the fold change matrix.

#### 2.3.2 Options for Adjusting Read Counts

Users can optionally provide multiplicative library size factors for each replicate in their experiment, since the total number of reads (sequencing depth) is usually unequal across replicates in RNA-seq experiments (Mortazavi *et al.*, 2008). All counts for a replicate will be multiplied by the library size factor.

GC (guanine-cytosine) content is known to affect expression measurements for genomic features, and the effect varies from sample to sample (Hansen *et al.*, 2012; Risso *et al.*, 2011; Benjamini and Speed, 2012). *Polyester* includes an option to model this GC bias in the simulated reads: for each biological replicate in the simulated data set, the user can choose one of 7 built-in GC content bias models, where one model was estimated from each of the 7 GEUVADIS replicates described in Section 2.2. We calculated these models using all transcripts from the available GEUVADIS transcriptome assembly (also described in Section 2.2).

For each replicate, we first calculated transcript-level read counts based on transcript length, sequencing depth, and the observed FPKM for the transcript. By definition of FPKM, read counts can be directly calculated using these inputs. We then centered the transcript counts around the overall mean transcript count, and modeled the centered counts as a smooth function of the transcript GC content using a loess smoother with span 0.3, analgous to smoothers previously used for modeling GC content (Benjamini and Speed, 2012).

Transcript GC content was calculated as the percentage of the annotated hg19 nucleotides falling in the boundaries of the assembled transcript that were G or C. The fitted loess curve defines a function that returns the average deviation from the overall mean transcript count for a transcript with a given GC content percentage. If there is no GC bias, the deviation would be 0. GC bias is added to replicates in *Polyester* after transcript-level counts have been specified by increasing or decreasing the count by the predicted deviation for that transcript’s GC content. The 7 loess curves included in *Polyester* are shown in Supplementary Figure 1. Users can also provide loess models from their own data as GC bias models if desired.

**Fig. 1.**
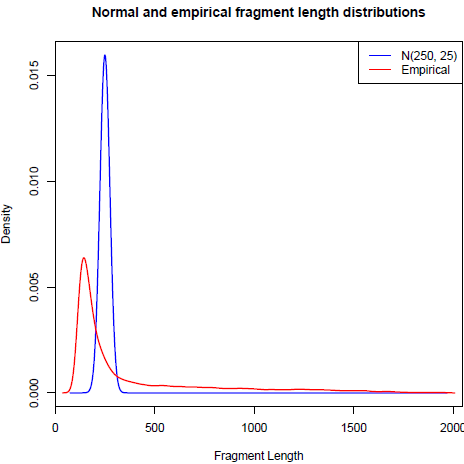
Fragment length distributions available in *Polyester*. The red curve shows the fragment length distribution for selected sequencing reads from the GEUVADIS RNA-seq data set; the blue curve shows a normal distribution with mean 250 and standard deviation 25. These two fragment length models are built into the simulator; users can also supply their own.

#### 2.3.3 User-Defined Count Models

As an alternative to the built-in negative binomial model, *Polyester* allows users to individually specify the number of reads to generate from each transcript, for each sample. This gives researchers the flexibility to design their own models for biological and technical variability, simulate complex experimental designs, such as timecourse experiments, and explore the effects of a wide variety of experimental parameters on differential expression results. This transcript-by-sample read count matrix can be created within R and input directly into *Polyester’s* read simulation function. This level of flexibility is not available with Flux Simulator or BEERS, which only allow specification of the total number of reads per replicate. While it is possible to write custom command-line scripts that induce differential expression using these simulators, differential expression models are built in to *Polyester*. This approach offers both a built-in model for convenience and an integrated way to define a custom model for flexibility.

### 2.4 The RNA Sequencing Process

#### 2.4.1 Fragmentation

After the transcripts have been specified and each transcript’s abundance in the simulated experiment has been set by determined by an assigned read count for each replicate, *Polyester* simulates the RNA sequencing process, described in detail in (Oshlack *et al*.,2010), beginning at the fragmentation step. All transcripts present in the experiment are broken into short fragments. There are two options for how fragment lengths are chosen: lengths can be drawn from a normal distribution with mean *μ_fl_* and standard deviation *σ_fl_* By default, μ_*fl*_ = 250 nucleotides and of*σ_fl_* = 25, but these parameters can be changed. Alternatively, fragment lengths can be drawn from an empirical length distribution included with the *Polyester* R package. This empirical distribution (Figure 1) was estimated from the insert sizes of the paired-end read alignments of the 7 GEUVADIS replicates described in Section 2.2, using Picard’s `CollectlnsertSizeMetrics` tool (BroadInstitute, 2014). The empirical density was estimated using the `logspline` function in R (Kooperberg and Stone, 1992; Kooperberg, 2013). Users can also supply their own fragment length distribution in `logspline` format. This distribution may be estimated from a user’s data set or varied to measure the effect of fragment length distribution on downstream results.

Ideally, the fragments generated from a transcript present in the sequencing sample would be uniformly distributed across the transcript. However, coverage across a transcript has been shown to be non-uniform (Mortazavi *et al*.,2008; Lahens*et al*., 2014; Li and Jiang, 2012). In *Polyester*, users can choose to generate fragments uniformly from transcripts, or they can select one of two possible positional bias models. These models were derived by Li and Jiang (2012), and they were based on two different fragmentation protocols.

The first model is based on a cDNA fragmentation protocol, and reads are more likely to come from the 3’ end of the transcript being sequenced. The second model incorporates bias caused by a protocol relying on RNA fragmentation, where the middle of each transcript is more likely to be sequenced. Both these models were estimated from Illumina data. Since the exact data from Li and Jiang (2012) was not made available with the manuscript, we extracted the data from Supplementary Figure S3 of Li and Jiang (2012) ourselves, using WebPlotDigitizer (Rohatgi, 2014), which can estimate the coordinates of data points on a scatterplot given only an image of that scatterplot. For reference, the figure is reproduced here (Supplementary Figure 2), created using the probabilities included as data sets (`cdnaf.rda` and `rnaf.rda`) in the *Polyester* R package.

**Fig. 2.**
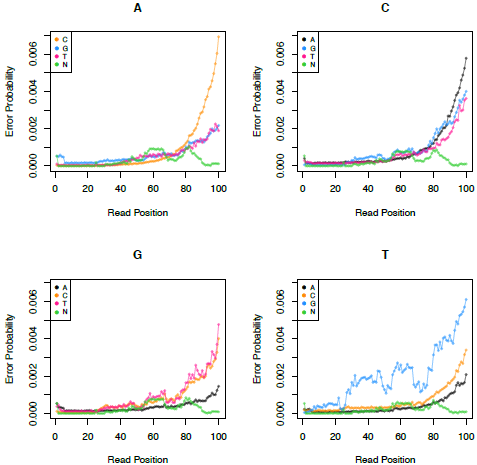
Example error model available in *Polyester*. Empirical error model derived from TruSeq SBS Kit v5-GA chemistry, using Illumina Genome Analyzer IIx, for mate 1 of a paired-end read. Separate panels are shown for each possible true reference nucleotide. Each panel illustrates the probability (y-axis) of mis-sequencing that reference nucleotide in a given read position (x-axis) as any of the 3 other nucleotides, or as an “N” (indicating an “unknown” nucleotide in the read). As expected, error probabilities increase toward the end of the read. Other error models, including the model for mate 2 of the read on this protocol, are illustrated in Supplementary Figures 3–7. If these error models are not suitable, custom error models can be estimated from any set of aligned sequencing reads.

#### 2.4.2 Sequencing

*Polyester* simulates unstranded RNA-seq reads in a manner compatible with the Illumina paired-end protocol (Sengupta *et al.*, 2011). In this protocol, read sequences are read off of double-stranded cDNA created from mRNA fragments, separated from other types of RNA using poly-A selection. To mimic this process in *Polyester*, each fragment selected from an original input transcript is reverse-complemented with probability 0.5: this means the read (for single-end experiments) or mate 1 of the read (for paired-end experiments) is equally likely to have originated from the transcript sequence itself and from the cDNA strand matched to the transcript fragment during sequencing.

Reads are then generated based on these fragments. A single-end read consists of the first *R* nucleotides of the fragment. For paired-end reads, these first *R* nucleotides become mate 1, and the last *R* nucleotides are read off and reverse-complemented to become mate 2. The reverse complementing happens because if mate 1 came from the actual transcript, mate 2 will be read from the complementary cDNA, and if mate 1 came from the complementary cDNA, mate 2 will come from the transcript itself (Illumina, 2011). By default, *R* =100 and can be adjusted by the user.

Users can choose from a variety of sequencing error models. The simplest one is a uniform error model, where each nucleotide in a read has the same probability *p_e_* of being sequenced incorrectly, and every possible sequencing error is equally likely (for example, if there is an error at a nucleotide which was supposed to be a T, the incorrect base is equally likely to be a G, C, A, or N). In the uniform error model, *p_e_ =* 0.005 by default and can be adjusted.

Several empirical error models are also available in *Polyester*. These models are based on two dataset-specific models that ship with the GemSim software(McElroy *et al.*, 2012). Separate models are available for a singleendread, mate 1 of a pair, and mate 2 of a pair, from two different sequencing protocols:Illumina Sequencing Kit v4 and TruSeq SBS Kit v5-GA (both from data sequenced on an Illumina Genome Analyzer IIx). These empirical error models include estimated probabilities of making each of the 4 possible sequencing errors at each position in the read. In general, empirical error probabilities increase toward the end of the read, and mate2 has higher error probabilities than mate 1 of a pair, and the TruSeq SBS Kit v5-GA error probabilities were lower than the Illumina Sequencing Kit v4 error probabilities(Figure 2; Supplementary Figure 3–7

**Fig. 3.**
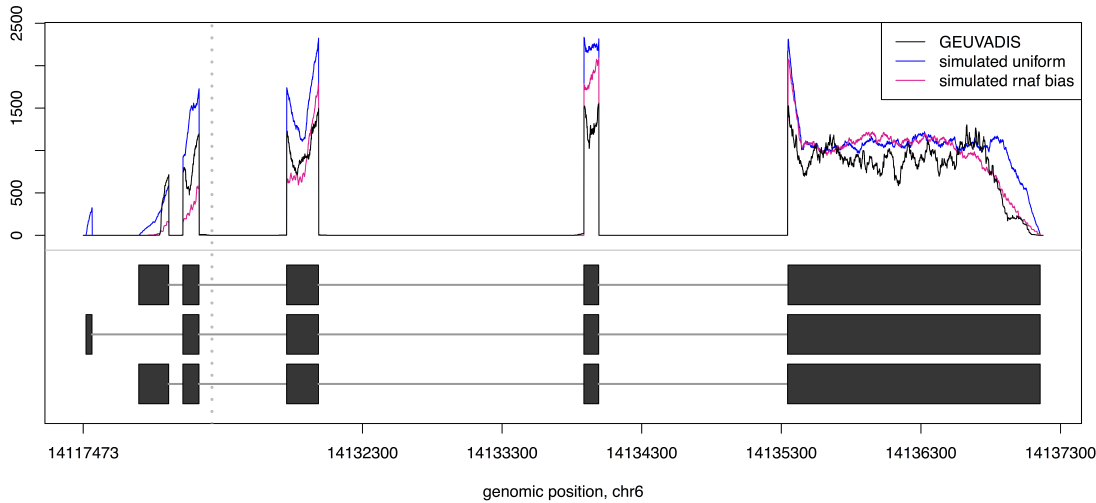
Coverage comparison to GEUVADIS data set. We counted the number of reads estimated to have originated from each of these annotated transcripts from gene CD83 (bottom half of figure) in the GEUVADIS RNA-seq data set, then simulated that same number of reads from each transcript using Polyester and processed those simulated reads. This figure shows the coverage track (y-axis, indicating number of reads with alignments overlapping the specified genomic position) for sample NA06985 (black), reads simulated without positional bias (blue), and read simulated using the rnaf bias model (pink). While the simulated coverage tracks look a bit cleaner than the track from the GEUVADIS data set, many of the major within-exon coverage patterns are captured in the simulation, especially with the uniform model. For example, both simulations capture the eak at the beginning of the rightmost exon. Note: the gray dotted line indicates that part of a long intron at that location was not illustrated in this plot.

*Polyester* can also handle custom error models: users can estimate an error model from their own sequencing data with the GemErr utility in GemSim. Detailed instructions on how to do this in a way compatible with *Polyester*are available in the package vignette.

After generating sequencing reads and simulating sequencing error, reads are written to disk in FASTA format. The read identifier in the FASTA files specifies the transcript of origin for each read, facilitating assessment of downstream alignment accuracy.Other pertinent simulation information is also automatically written to disk for use in downstream analysis: for each transcript, the transcript name, differential expression status, and fold change is recorded. For each replicate, the file name, group identifier *j*, and library size factor is recorded.

## 3 Results

### 3.1 Comparison with Real Data

To show that reads generated with *Polyester* exhibit realistic properties, we performed a small simulation experiment based on data from the 7 GEUVADIS RNA-seq replicates described in Section 2.2. For the experiment, we randomly selected 10 annotated genes with at least one highly-expressed isoform. We relied on the data-driven Cufflinks assembly to determine isoform expression: an annotated gene was considered to have highly-expressed isoforms if at least one of its annotated isoforms overlapped an assembled transcript with an average per-base coverage of at least 20 reads.

The 10 genes that were randomly selected had 15 transcripts between them: two had 3 isoforms, one had 2 isoforms, and the rest had 1 isoform. For the 10 genes, we counted the number of reads overlapping them using the `summarizeOverlaps` function in the Bioconductor package *GenomicAlignments* (Lawrence *et al.*, 2013). Counts were calculated from the TopHat-aligned reads from the GEUVADIS study for the 7 replicates described in Section 2.2. We then separated gene counts into isoform-level counts: we calculated per-isoform FPKM values for each of the 15 annotated transcripts using Cufflinks (Trapnell *et al.*, 2010) in its abundance-estimation-only mode, and used the FPKM ratio between isoforms of the same gene to generate isoform-level counts to simulate based on the gene counts we had already obtained.

We then used these isoform-level counts as input to *Polyester*, simulating a 7-replicate experiment with the specified number of reads being generated from each of the 15 selected annotated transcripts. Two experiments were simulated: one with all default options (no GC or positional bias, normal fragment length distribution with mean 250 and standard deviation 25, and uniform error model with 0.5% error probability) and one with all default options except for the positional bias model, for which we specified the rnaf bias model (Figure ??, red line).

The simulated reads were aligned to the hg19 genome with TopHat, and the coverage track for each experiment, for each simulated replicate was compared to the coverage track from the GEUVADIS replicate that generated the simulated replicate’s read count. For most of the transcripts, coverage tracks for both experiments looked reasonably similar to the observed coverage track in the GEUVADIS data set (see Figure 3 for a representative example).

The simulated coverage tracks were smoother than the coverage track from the GEUVADIS data set, but major trends in the coverage patterns within exons were captured by the simulated reads. There are annotated transcripts for which reads generated by *Polyester* do not adequately capture the observed coverage in the GEUVADIS data set (Supplementary Figure 8, especially when positional bias is added. This seems to mainly occur in cases where only a very small part of a large exon appears to be expressed in the data set (as is the case in Supplementary Figure 8. The coverage for most of the other transcripts was similar to the real data for most genes and replicates (Frazee, 2014). Reads simulated with rnaf bias sometimes had poor coverage for genes consisting of transcripts with many small exons.

For these 15 simulated transcripts, FPKM estimates were positively correlated between each simulated data set and the GEUVADIS data set for each replicate. To get data for this comparison, we used Cufflinks’s abundance-estimation-only mode to get expression estimates for the 15 isoforms based on the simulated reads’ alignments, in the same way we calculated expression for the GEUVADIS replicates. We calculated correlation between FPKM estimates of the 15 transcripts for the GEUVADIS data set and for each of the simulated data sets, using correlation instead of absolute FPKM because normalization for number of mapped reads put the sets of FPKMs on different scales.

For the simulation without positional bias, the correlation was extremely high: the minimum correlation across the 7 replicates studied was 0.98. However, the FPKM estimates were less correlated when RNA-fragmentation-related positional bias was induced: all correlations were positive, but weak (Supplementary Figure 9). These results generally indicate that realistic coverage profiles can be obtained with *Polyester* but that adding positional bias may cause problems when transcripts have unusual structure. The correlation in FPKM estimates between the simulated data sets and the GEUVADIS samples suggests that *Polyester* captures transcript level variation in gene expression data.

### 3.2 Use Case: Assessing the Accuracy of a Differential Expression Method

To demonstrate a use case for *polyester*, we simulated two small differential expression experiments and attempted to discover the simulated differentialexpression using *limma* (Smyth, 2005).

The first experiment used the default size parameter in *Polyester*,which means the variance of the distribution from which each transcript’s count is drawn is equal to 4 times the mean of that distribution. In other words, the mean and variance of the transcript counts are linearly related. We refer to this experiment as “low variance.” The second experiment set the size parameter to 1 forall transcripts, regardless of the mean count, which means each transcript’s mean and variance are quadratically related. This experiment was the “high variance” experiment.

In both scenarios, the main wrapper function in *Polyester* was used to simulate classic two-group experiments. Reads were simulated from transcripts on human chromosome 22 (hg19 build, *N* = 926).*μ_k_* was set to length(transcript*_k_*)/5, which correspondsto approximately 20x coverage for reads of length 100. We randomly chose 75 transcripts to have λ = 3 and 75 to have λ = 1/3; the rest had λ =1. For *n_j_ =* 7 replicates in each group *j*, we simulated paired-end reads from 250-base fragments (σ*_fl_* = 25), with a uniform error probability and the default error rate of 0.005. Simulated reads were aligned to hg19 with TopHat 2.0.13 (Trapnell *et al.*, 2009), and Cufflinks 2.2.1 (Trapnell *et al.*, 2010) was used to obtain expression estimates for the 926 transcripts from which transcripts were simulated. Expression was measured using FPKM (fragments per kilobase per million mapped reads). We then ran transcript-level differential expression tests using *limma* (Smyth, 2005). Specifically, for each transcript k, the following linear model was fit: 
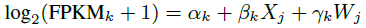
where FPKM*_k_* is the expression measurement for transcript *k*, *X_j_* is 0 or 1 depending on which group sample *j* was assigned to, and *W_j_* is a library-size adjustment, defined as the 75th percentile over all *k* of the log2 (FPKM*k* + 1) values for replicate *j* (Paulson *et al.*,2013). We fit these linear models for each transcript, and for each β*_k_*, we calculated moderated *t*-statistics and associated p-values using the shrinkage methodology in *limma*’s eBayes function. We calculated ROC curves based on these p-values and our knowledge of the true differential expression status of each transcript. Sensitivity and specificity of the *limma* differential expression analysis were high for the small-variance scenario, but were diminished in the large-variance scenario, as expected (Figure 4).

**Fig. 4.**
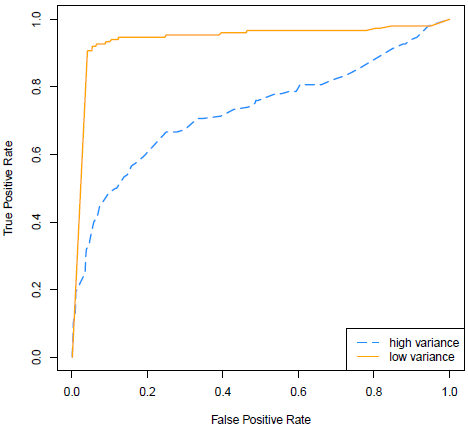
ROC curves for transcript-level differential expression calls from *Polyester* data sets. For varying significance (p– or q-value) cutoffs,sensitivity and specificity from the simulation experiments. Differential expression was more difficult to detect under conditions where expression levels were highly variable between replicates,as expected.

Since expression fold changes can be explicitly specified in *Polyester*, we can also investigate whether those fold changes are preserved throughout this RNA-seq data analysis pipeline (Figure 5). In general, the coefficient distributions for transcripts not specified to be differentially expressed were centered around zero, as expected, since models were fit on the logscale. The coefficient distributions should have been centered around log_2_ (3) = 1.58 for the overexpressed transcripts (expression level three times higher in the first group), and around loglog_2_(1/3) = −1.58 for the underexpressed transcripts (expression level three times higher in the second group). The overexpressed distributions had means 1.39 and 1.44 in the high- and low-variance scenarios, respectively, and the underexpressed distributions had means −1.57 and −1.60 in the high– and low-variance scenarios, respectively. Coefficient estimates were much more variable in the scenario with higher expression variance (Figure 5). These numbers are similar to the the specified value of 1.58, indicating that the RNA-seq pipeline used to analyze these data sets satisfactorily captured the existence and magnitude of the differential expression set in the experiment simulated with *Polyester*.

For this differential experiment, where about 639,000 reads per sample were simulated, read generation took 1–2 minutes per biological replicate in the experiment and 4.4G memory was used on a single cluster node with one core.

**Fig. 5.**
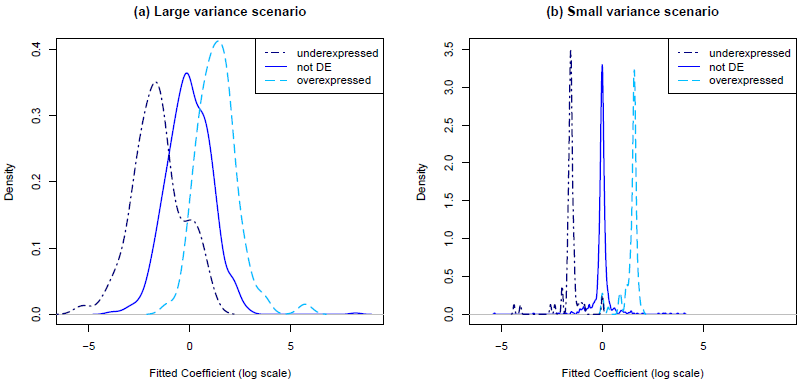
Coefficient distributions from differential expression models. Distributions from the high-variance scenario are shown in panel (a) and from the low-variance scenario are shown in panel (b). These distributions of estimated log fold changes between the two simulation groups tend to be centered around the values specified at the beginning of the simulation, and there is more variability in the coefficient estimates for high-variance scenario, as expected.

These examples illustrate some of the many possible ways *Polyester*can be used to explore the effects of analysis choices on downstream differential expression results.

## 4 Discussion

In this paper, we propose a lightweight, flexible RNA-seq read simulator allowing users to set differential expression levels at the isoform level. A full experiment with biological replicates can be simulated with one command, and time-consuming alignment is not required beforehand.

The sequencing process is complex, and some subtleties and potential biases present in that process are not yet implemented in *Polyester* but could be in the future. For example, adding random hexamer priming bias (Hansen *et al.*, 2010), implementing PCR amplification bias (Fang and Cui, 2011) or other biases that depend on the specific nucleotides being sequenced, simulating quality scores for base calls, and adding the ability to simulate indels are all possibilities for future improvements. However, our comparisons with real data suggest that the *Polyester* model sufficiently mimicks real sequencing data to be practically useful.

## 5 Software

Polyester is available from Bioconductor: http://bioconductor.org/packages/release/bioc/html/polyester.html. The development version is available on GitHub:https://github.com/alyssafrazee/polyester.Community contributions and bug reports are welcomed in the development version. Code for the analysis shown in this paper is available at https://github.com/alyssafrazee/polyester_code.

## Acknowledgements

*Funding*: JL and BL are supported by NIH R01 GM105705. AF is supported by a Hopkins Sommer Scholarship. AJ is supported by the Lieber Institute for Brain Development.

